# Brain-wide visual habituation networks in wild type and *fmr1* zebrafish

**DOI:** 10.1101/722074

**Authors:** Emmanuel Marquez-Legorreta, Lena Constantin, Marielle Piber, Itia A. Favre-Bulle, Michael A. Taylor, Gilles C. Vanwalleghem, Ethan Scott

## Abstract

Habituation is a form of learning during which animals stop responding to repetitive stimuli, and deficits in habituation are characteristics of several psychiatric disorders. Due to the technical challenges of measuring brain activity comprehensively and at cellular resolution, the brain-wide networks mediating habituation are poorly understood. Here we report brain-wide calcium imaging during visual learning in larval zebrafish as they habituate to repeated threatening loom stimuli. We show that different functional categories of loom-sensitive neurons are located in characteristic locations throughout the brain, and that both the functional properties of their networks and the resulting behavior can be modulated by stimulus saliency and timing. Using graph theory, we identify a principally visual circuit that habituates minimally, a moderately habituating midbrain population proposed to mediate the sensorimotor transformation, and downstream circuit elements responsible for higher order representations and the delivery of behavior. Zebrafish larvae carrying a mutation in the fmr1 gene have a systematic shift towards sustained premotor activity in this network, and show slower behavioral habituation. This represents the first description of a visual learning network across the brain at cellular resolution, and provides insights into the circuit-level changes that may occur in people with Fragile X syndrome and related psychiatric conditions.

---

Habituation is a simple form of non-associative learning, characterized by a decrease in response after multiple presentations of a stimulus, that is conserved across much of the animal kingdom^1^. It allows animals to remain attentive to novel and ecologically relevant stimuli while minimizing their expenditure of energy on inputs that occur frequently without consequence. The strength and speed of habituation, and of recovery during periods without the stimulus, depend on the parameters of the stimulus and its repetitions (the intensity, frequency, and number of stimuli)^2,3^. Careful modulations of these stimulus properties have proven useful in exploring the relationships between repetitive stimuli and behavior, thereby providing clues about the underlying habituation circuitry^4–7^.

Other work has addressed some of the molecular and cellular dynamics mediating habituation, including reductions in motor neurons’ presynaptic vesicle release during short-term habituation and processes involving protein syntheses for longer-term forms of habituation^8–13^. At the other end of the spectrum, fMRI studies in humans have revealed changes in activity for various brain regions during habituation^14–16^. The intervening scales, of regional circuits and brain-wide networks, cannot be addressed using targeted cellular techniques or traditional brain-wide approaches. These networks, and the ways in which they change during habituation, can only be addressed by observing activity in whole populations of neurons (up to and including the whole brain) at single-cell resolution.

In recent years, exactly this approach has become possible in zebrafish larvae through the use of genetically encoded calcium indicators and light-sheet or 2-photon microscopy^17^. Since zebrafish larvae undergo behavioral habituation^18,19^ they have been used for experiments with tactile, visual, and acoustic stimuli, exploring the genetic and molecular mechanisms of specific circuits^20–24^. Furthermore, they share important molecular underpinnings of habituation with other species^25–27^. All together, these features make them an appealing platform for exploring brain-wide habituation circuitry.

This approach requires a robust innate behavior that is subject to habituation. Looming visual stimuli, which simulate approaching predators, reliably elicit startle responses that are conserved from insects to humans^28^, and repeated looms have been shown to produce habituation in various species^29–31^. When looming stimuli are presented to larval zebrafish, visual information converges in the tectum, where local circuits are proposed to calculate the imminence of a threat^31–33^. However, additional structures respond to looms^31–35^, and others, including the hypothalamus, modulate the visual escape behavior in contexts other than habituation^34,36,37^. The result is an intriguing but sketchy outline of the habituation network, and in the absence of a whole-brain cellular-resolution analysis, numerous questions about this behaviorally important process remain unanswered.

Addressing these questions is especially important because of the role that sensorimotor transformations and habituation play in psychiatric disorders including schizophrenia, autism spectrum disorder (ASD), and Fragile X syndrome (FXS)^38^. While these disorders are traditionally diagnosed around their social or cognitive symptoms, each has characteristic alterations in sensory processing, habituation, and sensorimotor gating that compound, or in some cases may drive, social and intellectual impairments^39,40^. FXS patients, for example, show slow habituation^41–44^, a phenotype also found in *fmr1-*mutant mice that model FXS^45^. While fMRI and EEG studies have revealed some of the regional changes in neural activity that correlate with habituation deficits in various psychiatric disorders^46–49^, the network-wide causes of these symptoms remain largely unexplored.

## Habituation of visual escape behavior in larval zebrafish

To characterize the escape behavior of larval zebrafish exposed to looming stimuli, we designed a 12-well apparatus in which each well contained a larva receiving its own loom stimulus from below (Figure 1a). We presented looms in blocks of 10, with five minutes between blocks and an auditory tone at the end of the second rest period (for dishabituation before the 21^st^ loom stimulus). In order to explore the relationships between stimulus properties and behavioral habituation, we used looming stimuli of two expansion speeds (a fast stimulus that filled the bottom of the well in 2 sec and a slow stimulus taking 4 sec) and two inter-stimulus intervals (ISIs) of 20 or 60 sec between looms. This resulted in four stimulus trains: f20, f60, s20, and s60 (Figure 1b).

**Figure 1.**
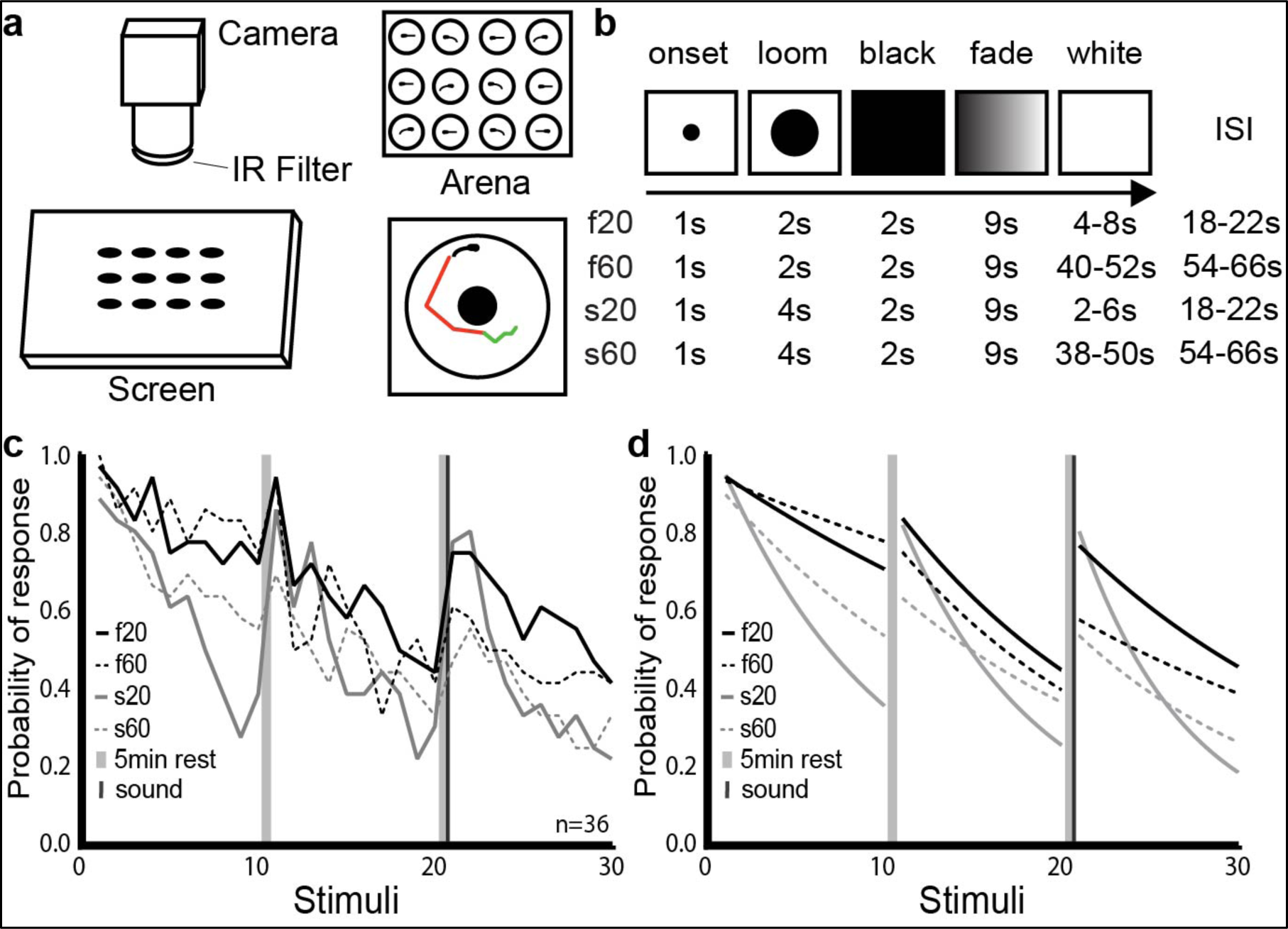
Modulation of habituation by stimulus features. **a**. Schematic representation of our setup for measuring visual habituation behavior. A 12-well chamber with one larva in each well (top right) was filmed on a horizontal screen (left) on which the looms were presented. Automated tracking recorded periods of swim bouts (green) and burst swim (red) for each larva (bottom right) **b**. Stimulus train properties across the 4 experimental groups. **c**. Probability of response across the 4 groups during three blocks of ten loom presentations. **d.** Smoothed curves of the response probability for each group.

Each led to habituation of loom-elicited startle responses (Figure 1c), and two patterns arose across the four stimulus trains. First, the slow-growing stimuli led to stronger habituation than the fast stimuli did, especially in the first block of 10 looms. Second, the stimulus trains with 20 sec ISIs produced faster habituation within blocks, but the habituation produced by trains with 60 sec ISIs showed less recovery after the 5-minute rest periods. A generalized linear mixed model (GLMM) of the first block indicated a significant effect of the loom presentation number (beta= −0.25365, p= 2.00 × 10^−16^) on response probability, confirming habituation. The loom speed also affected response probability strongly (beta= −1.23839, p= 2.22 × 10^−8^) with a weaker but significant impact from the ISI (beta= 0.45089, p= 0.038). Together, the speed, ISI, and presentation number explain almost 20% of the variance (R^2^= 0.1864) and together with the random variable (fish identity) the model explained more than 35% of the variance (R^2^= 0.3647). These effects are consistent with past studies in zebrafish and other diverse model systems^5,7,26,27^, suggesting a relationship between stimuli and habituation behavior that is broadly conserved. Explaining this relationship requires an exploration of the underlying circuitry and the ways in which it changes during habituation.

## Brain-wide characterization of neural activity during habituation

To address brain-wide patterns of activity during habituation, and the types of individual neurons that drive them, we moved to a head-embedded preparation in which loom stimuli were presented on an LCD screen. We performed whole-brain imaging of the *elavl3:H2B-GCaMP6s* line using selective plane illumination microscopy (SPIM), as previously described (See Online Methods). For each larva, this produced 50 dorso-ventral planes, at 5µm intervals, spanning the rostro-caudal and medio-lateral extents of the brain, with a volumetric acquisition rate of 2Hz. We performed morphological segmentation of these images to identify regions of interest (ROIs) generally corresponding to individual neurons, and extracted fluorescent traces from these ROIs, as described before (See Online Methods).

Snapshots of responses across the brain during this repetitive stimulation (shown for f20 in Figure 2a-c) show a sharp decrease in responsive ROIs between the first and second stimuli, and a further drop in responses by the 10^th^ stimulus. Figure 2d and 2e show the response of each ROI in the second and 10^th^ trial as a proportion of its response in the first. Habituation is conspicuous across all loom-responsive brain regions, including the tectum, thalamus, medial hindbrain, tegmentum, and telencephalon, suggesting that these regions are affected by or involved in the habituation process.

**Figure 2.**
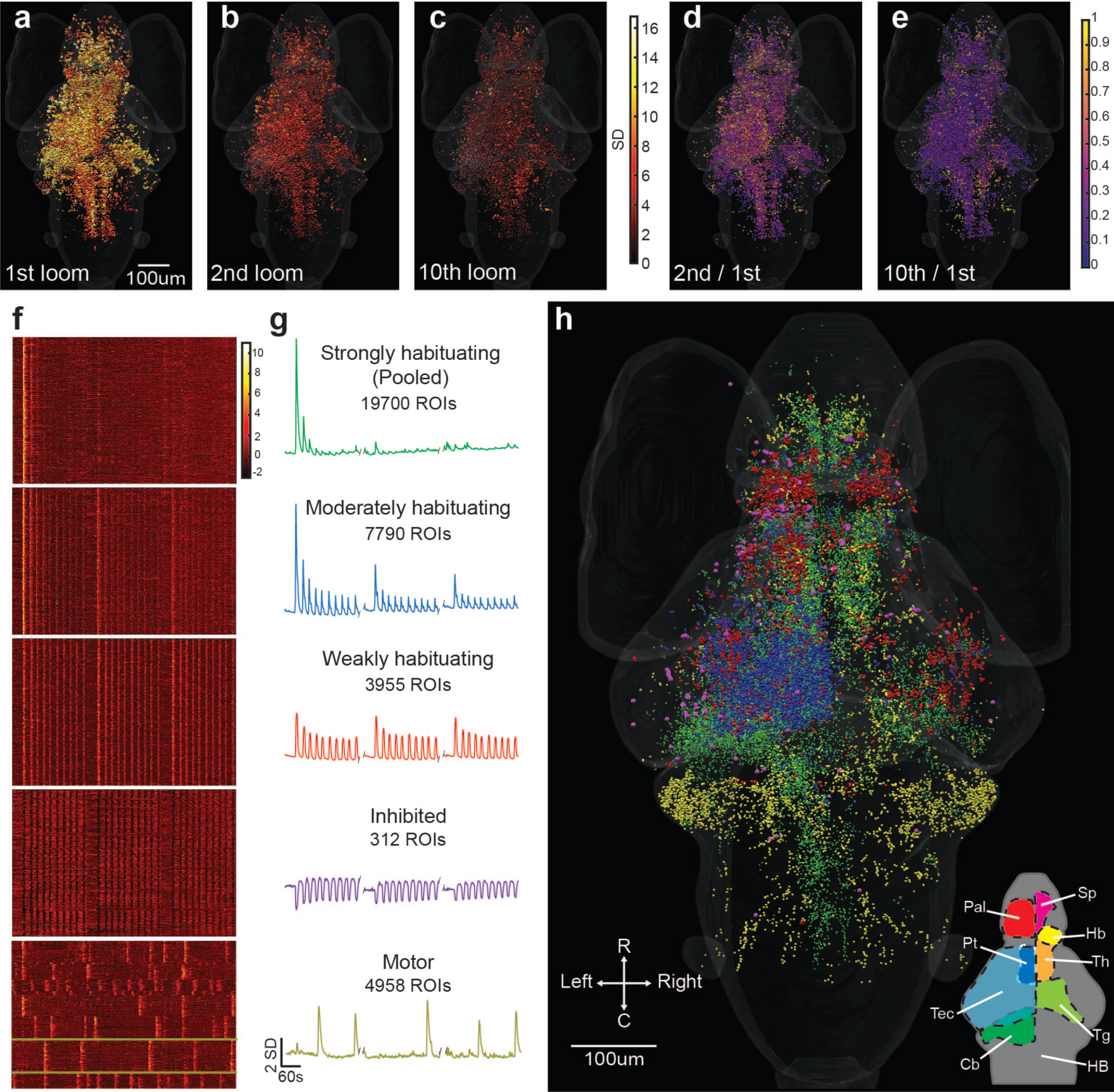
Activity of individual ROIs and their functional clusters during habituation. **a.** Responses of ROIs across the brain to a loom stimulus, color coded for the intensity of their response. **b, c**. The same ROIs’ responses to the second and tenth looms. **d, e**. The degree of habituation in each of these ROIs in the second and 10^th^ trials, calculated as the ratio of response to the first loom. This analysis was restricted to ROIs showing clear responses (with a coefficient of determination (r^2^ value) >0.5 for the linear regression between their response and a regressor simulating a calcium signal) for the first loom stimulus. Raster plots (**f**) and mean responses **(g)** of the ROIs composing each of five functional clusters, with a clear correspondence to the three blocks of ten stimuli. **h.** Anatomical locations for the ROIs belonging to each functional cluster. Since different animals startled in different trials, we identified the motor cluster using a different regressor for each animal. The mean responses are shown for a single animal in this cluster in **g**, with yellow lines indicating the relevant neurons from that animal in **f**. A rotation of **h** can be found in Supplementary Movie 1, and the distributions of these clusters is detailed in virtual sections in Supplemental Figure 3. Data show the pooled responses of 11 larvae to the f20 stimulus train. Relevant anatomical brain regions are indicated in the bottom right corner of **(h)**, each shown for only one side of the brain. Pallium, Pal; subpallium, Sp; thalamus, Th; habenula, Hb; pretectum, Pt; tectum, Tec; tegmentum, Tg; cerebellum, Cb; and hindbrain, HB. R, rostral; C, caudal.

To address these possible mechanisms, we used k-means clustering to identify seven categories (clusters) of loom-responsive neurons with distinct functional properties (and one auditory cluster, not shown as there was no significant dishabituation). Based on their highly similar response properties, we merged three clusters of ROIs showing strong and rapid habituation (Supplemental Figure 1) into a single strongly habituating cluster (Figure 2f, g). We characterized the remaining three clusters as moderately habituating, weakly habituating, inhibited, and we also located a motor-associated group of ROIs using regressors customized to each animal’s movements (Figure 2f, g). A t-SNE analysis (Supplemental Figure 2) shows functional segregation among these clusters, suggesting that they are distinct categories of loom-responsive neurons.

Strongly habituating ROIs are spread across several brain regions (Figure 2h and Supplemental Figure 3), most prominently in the tectum, thalamus, medial hindbrain, pallium, and tegmentum. In the hindbrain, these ROIs are concentrated in a longitudinal rostro-caudal strip along the pathway of the tectobulbar projections, meaning that they likely include reticulospinal premotor neurons^50,51^.

**Figure 3.**
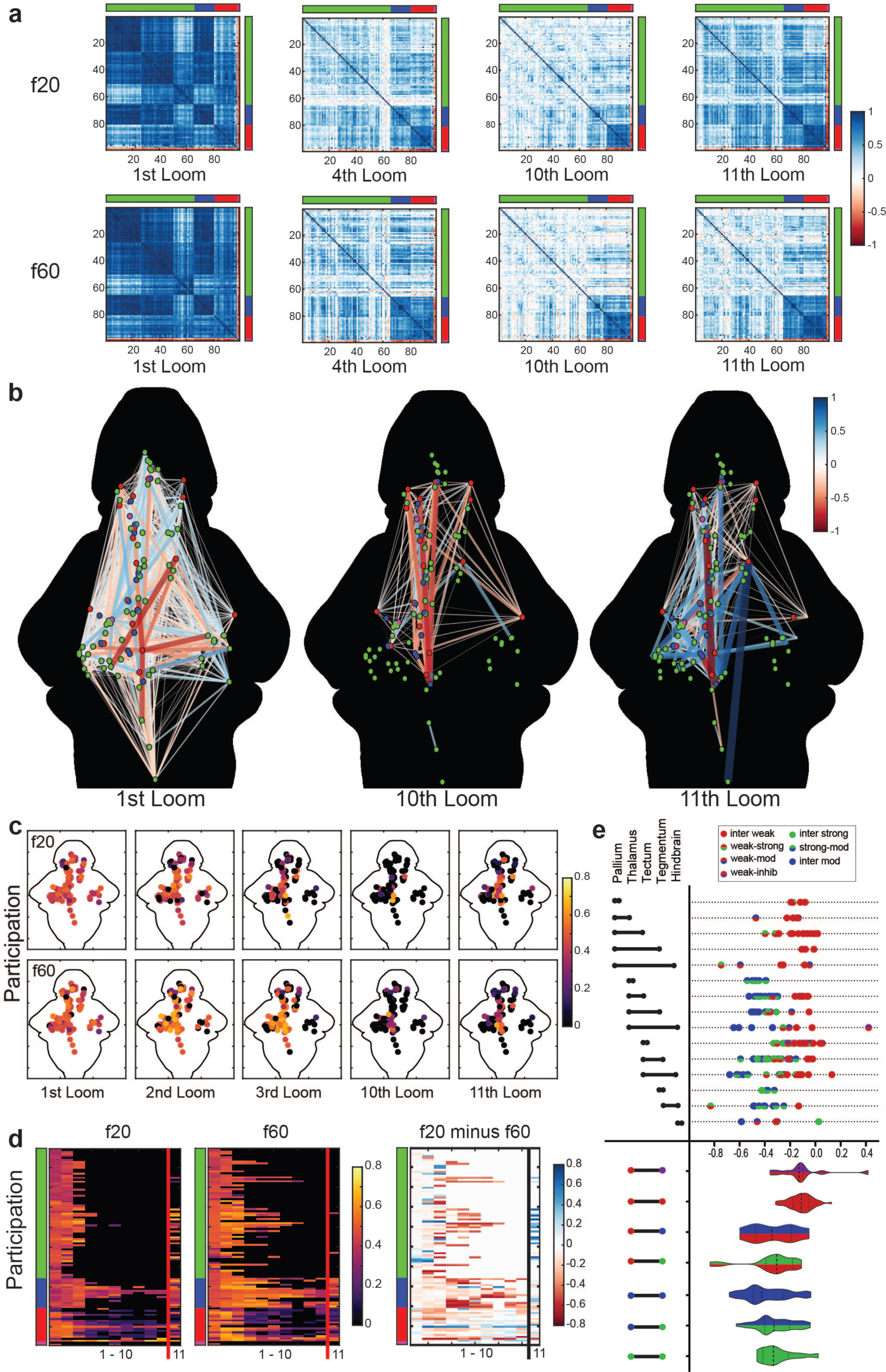
The visual loom network, and the changes that occur during habituation. **a.** Correlation matrices for activity across 99 nodes representing ROIs across the whole brain. The functional clusters to which each node belongs are indicated on the axes, using the color code from Figure 2. Darker blue shades represent stronger positive correlations for any given pairing, and red indicates negative correlations (see color scale, **a**). **b**. A graphic representation of correlations across the 99 nodes, whose functional clusters are indicated by their colors and anatomical positions represented spatially. The colors and width of the lines indicate the relative correlation across the f20 and f60 experiments (f20 correlation minus f60 correlation), where red indicates stronger correlations in f60 and blue indicates stronger correlations in f20 (see color scale). Only edges with correlations above 0.75 in either the f20 or the f60 matrices are displayed. **c.** A heat map of the participation for each of the 99 nodes during the 1^st^, 2^nd^, 3^rd^, 10^th^, and 11^th^ loom stimuli of the f20 and f60 experiments. **d.** Raster plots showing the participation of each node across the first 11 stimuli for f20 and f60, and the relative participation (f20 value minus f60 value) where blue indicates stronger f20 participation and red indicates stronger f60 participation. The functional clusters for each node are indicated, using the color code from Figure 2. **e.** Changes in correlation strength for all inter-node edges from the 10^th^ to the 11^th^ looms of f20, indicating the impacts of the recovery from habituation. Values shown are calculated for each edge as its correlation in the 10^th^ loom minus its value in the 11^th^ loom, with more negative values indicating edges that showed more pronounced recovery between the 10^th^ and 11^th^ looms (top). The functional clusters for each edge’s two nodes are color coded and the brain regions that the edges span are indicated on the left. Violin plots (bottom) show the cumulative distributions of edges connecting different types of functional clusters, as indicated on the left.

Moderately habituating ROIs are tightly concentrated in the central region of the tectal periventricular layer (PVL) of the left tectum (Figure 2h and Supplemental Figure 3). This laterality is unsurprising, since the stimulus was presented to the right eye, and since all retinal projections are contralateral in zebrafish larvae. This position is consistent with a role for the associated neurons in the spatially registered processing of visual information, and their decreased responses may represent an important element of the overall circuit’s reduced responsiveness during habituation.

Weakly habituating ROIs are prominent in the tectum, habenulae, pretectum, and pallium (Figure 2h and Supplemental Figure 3). There is moderate laterality toward the contralateral side to the stimulus in most of these regions. In the pallium, responses are concentrated around the dorsal edge of the pallium in what will likely become the lateral division of the dorsal pallium (Dl), although they also extend into the medial division (Dm, Supplemental Figure 3).

Inhibited ROIs are rare and mostly localized to the contralateral tectum and rostral thalamus (Figure 2h and Supplemental Figure 3). Motor-associated ROIs are concentrated in the cerebellum. However, some can be found in the anterior and lateral hindbrain and small numbers occur in the thalamus and pallium (Figure 2h and Supplemental Figure 3). These ROIs are presumably involved in the coordination and delivery of the escape responses.

## Temporal stimulus properties modulate brain-wide responses

By looking at the changes in these distributions across our four stimulus trains (Figure 1), we next aimed to characterize brain-wide activity under conditions that lead to different rates and persistence of habituation in free-swimming larvae. The fundamental brain-wide habituation network was conserved across these treatments, but specific functional differences emerged (Supplemental Figure 4a). One was a greater number of strongly habituating ROIs in the hindbrain strip for the s20 and s60 experiments, with possible relevance to the faster habituation that slow stimuli drive (Figure 1). Another came in experiments with 60 sec ISIs, where we observed a greater number of weakly habituating ROIs in the dorsal hindbrain on the side contralateral to the stimulus (Supplemental Figure 4). This may play a role in the stronger preservation of habituation across breaks in the 60 sec ISI experiments.

**Figure 4.**
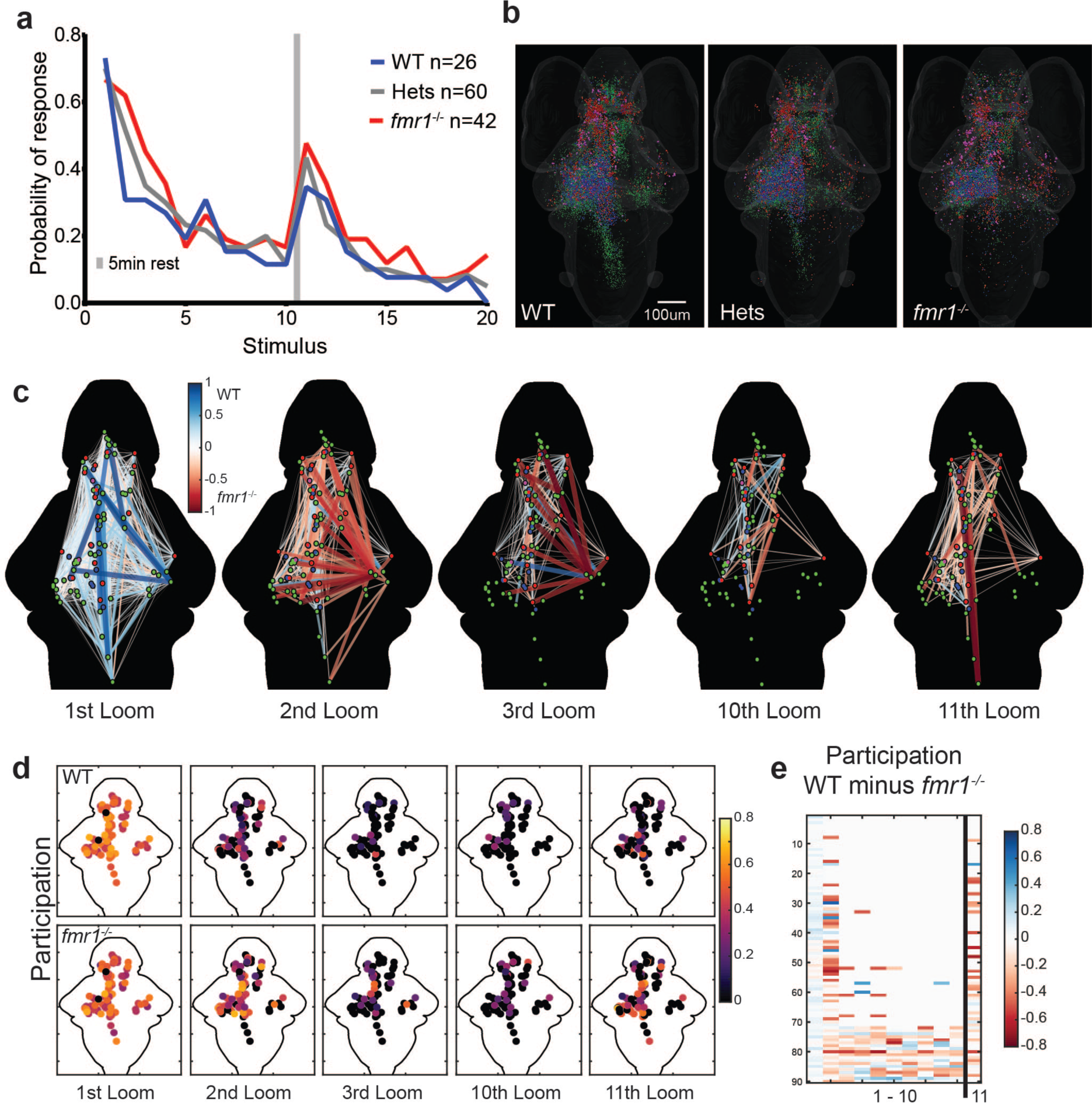
Behavioral and network-wide changes in fmr1^−/−^ larvae. **a**. Over the course of two blocks of 10 stimuli, *fmr1^−/−^* larvae show slower habituation and stronger recovery than WT siblings, and heterozygotes show an intermediate phenotype. Binomial test: *fmr1^−/−^* versus WT: 2^nd^ Loom (p=3.056e^−5^); 3^rd^ Loom (p=0.034); 11^th^ Loom (p=0.055) and 16^th^ Loom (p=0.039). Hets versus WT: 2^nd^ Loom (p=0.001) and 9^th^ Loom (p=0.039). All other comparisons (p>0.1). **b**. Brain-wide distributions of ROIs for the three genotypes, color coded for functional cluster as in Figure 2 and Supplemental Figure 4. **c**. Node-based graphs showing relative correlations (WT correlation minus *fmr1^−/−^* correlation), where blue indicates correlations that are stronger in WT and red indicates correlations that are stronger in *fmr1^−/−^*. **d**. Heat maps of participation for all nodes across habituation and recovery. **e.** A raster plot of relative participation (WT participation minus *fmr1^−/−^* participation) for each node through the first 11 trials.

Since a large proportion of loom-responsive ROIs are in the tectum, especially for the moderately habituating cluster (Supplemental Figure 4b), we next looked at the relationship between the stimulus train’s properties and the responses of each functional cluster in the tectum (Supplemental Figure 4c). This revealed only subtle difference across the stimulus trains for the response profiles of fast habituating neurons. For moderately habituating neurons, differences arose with intriguing parallels to the behavioral outputs. In experiments with 60sec ISIs, habitation is slower, and recovery is less dramatic than for 20sec ISI experiments. In experiments that used slow stimuli, habituation occurs faster than in the corresponding experiments with fast loom stimuli. For weakly habituating neurons, experiments with 60sec ISIs lead to less habituation throughout the experiment, while other correlates of behavior are less clear. Overall, moderately habituating ROIs repeatedly had the strongest correlation to free-swimming escape probability (Pearson correlation values: f20= 0.6896; f60= 0.6896; s20= 0.6368; s60= 0.7724), suggesting that among our functional clusters, it is the moderately habituating ROIs in the tectum whose dynamics most closely reflect behavior.

## Network Modelling of Visual Loom Habituation

As an approach to modelling visual loom processing and the network changes that produce habituation, we applied graph theory to our brain-wide activity data. To generate a tractable dataset, we downsampled our 144,709 responsive ROIs into 99 nodes that represent the ROIs’ functional clusters and anatomical locations, and then produced matrices representing the correlations in activity across each of these nodes at different times during the experiments (Figure 3a, see Online Methods). We then compared these matrices in larvae exposed to the f20 and f60 habituation paradigms to identify the network-level correlates of behavioral habituation. As expected, both paradigms produced high correlation values in response to the first loom, and the matrices for the two paradigms were highly similar. As habituation proceeded, network correlations remained somewhat higher in the f60 paradigm, reflecting differences in the behavioral responses during the f20 and f60 experiments (Figure 1 c, d). By the 10^th^ loom, most of these correlations had dropped dramatically for both paradigms, with high values mostly restricted to correlations between weakly habituating (red) nodes. The f20 paradigm shows a stronger recovery across the network in the 11^th^ trial, reflecting the stronger behavioral recovery that takes place in this paradigm.

As an approach to judge both the rate at which these correlations were lost during the first block of stimuli and the degree to which they recovered in the 11^th^ trial, we used a Pearson correlation to match the matrix of the 11^th^ trial to the most closely related matrix from the first block of stimuli. The highest Pearson correlation coefficients were for the 4^th^ trial for f20 and the 6^th^ trial for f60, indicating both that the correlations are lost more quickly in f20 (the paradigm in which habituation occurs more quickly), and that the recovery is weaker in f60 (the paradigm that produces more indelible behavioral habituation). Notably, the patterns of correlations across the matrices during mid-habituation trials (4^th^ for f20, Figure 3a, and 6^th^ for f60, now shown) strikingly resemble those in the 11^th^ trials, suggesting that the network is returning to a partially habituated state.

These results show that the loss of correlations across nodes in the network reflects behavioral outputs. To describe the networks containing these nodes, we represented them spatially and mapped the relative correlation strengths between nodes in the f20 and f60 paradigms (Figure 3b). Each edge (node-to-node relationship) in the graph is represented by its correlation value in the f20 paradigm minus its value in f60 paradigm. As expected, because the first trial is identical, both paradigms show robust correlations across numerous edges in the first trial, with most edges near a zero value and no net weighting of the graph toward positive or negative. By the 10^th^ trial, the graph has lost most edges, and the remaining activity is biased toward stronger correlations in f60 (shown in red), reflecting the slower habituation. The f20 paradigm shows stronger recovery, however, and this is captured in a shift toward positive values (blue) in the 11^th^ trial.

We then quantified the participation of each node in the graph, where participation is defined as the proportion of a node’s highly correlated edges that are shared with nodes from a different functional cluster (as defined in Figure 2). Participation dropped over the course of 10 stimuli (Figure 3c), but this drop was slower in f60, suggesting that habituation is driven not only by a drop in correlation across nodes, but specifically by a loss of communication between different functional clusters. This is reinforced by the higher participation in the 11^th^ trial of the f20 paradigm, where strong behavioral recovery is echoed by a recovery in participation. Raster plots of participation by each node across the first 11 trials (Figure 3d) show this trend, further suggesting that it is weakly habituating (red nodes) that maintain much of their participation as habituation proceeds, and that recovery is accompanied by a resumption of participation by various strongly (green) and moderately (blue) habituating nodes.

To address which brain regions are involved in this process, we mapped the correlation strengths of edges between nodes across five regions containing a majority of the nodes (the pallium, thalamus, tectum, tegmentum, and hindbrain, Figure 3e). The values for each edge, represented by a dot, show the correlation in the 10^th^ trial minus the correlation in the 11^th^ trial, thus giving negative values to edges that became stronger during recovery. Violin plots show the total distributions of edges between different functional clusters. The results confirm that certain types of edges, especially those between two weakly habituating (red) nodes, play a relatively small role in recovery, owing to their strong unhabituated responses in the 10^th^ trial. Other types of edges, especially those not including a weakly habituating node, tend to have highly negative values, indicating that they contribute to the part of the network that is lost during habituation and regained during recovery.

Collectively, these results converge to produce a model of the brain-wide network that produces visual escape and the mechanisms by which these responses are suppressed during learning. The initial process of habituation appears to rest on the loss of correlation (and presumed communication) among neurons of different functional clusters. This is manifested as a dramatic drop in correlation values for edges between different clusters (Figure 3a), the restriction of the active network principally to edges between nodes of the same type (especially weakly habituating nodes, Figure 3b, e), and the loss of participation during the course of habituation (Figure 3c). The striking similarity between mid-habituation matrices and those of partially recovered networks (Figure 3a) indicates that the matrix changes that underlie habituation are the same as those that are reversed during recovery, suggesting that the onset of habituation works through the same circuit-level changes that recovery does, and that there are not separable network-level mechanisms for the acquisition and retention of behavioral habituation.

## *fmr1*^−/−^ mutant larvae show behavioral and network-level habituation deficits

To test the validity and explore the utility of this proposed network, we next looked for phenotypes across the habituation network in a zebrafish model of FXS, an inherited disorder characterized by intellectual disability, social deficits, and sensory phenotypes. We used a nonsense mutation in the highly conserved *fmr1* gene, the perturbation of which causes FXS in humans. Given the learning deficits, including slow habituation^42–44^, in humans with FXS, we explored whether and how behavior and brain-wide habituation networks are altered in *fmr1*-mutant zebrafish.

Using the s20 habituation paradigm in our free-swimming preparation, we found that *fmr1^−/−^*, *fmr1^−/+^* heterozygotes (hets), and wild type (WT) siblings share a similarly high probability of startling to the first loom stimulus (Figure 4a). Habituation is slower, however, and recovery after a break is more dramatic, in *fmr1^−/−^* animals. Heterozygotes show an intermediate phenotype.

We next looked for correlates of this behavior using brain-wide calcium imaging, first looking at the distributions of ROIs belonging to functional clusters (Figure 4b). While all genotypes had fundamentally similar distributions, there was a trend toward more numerous weakly habituating ROIs in the cerebellum in *fmr1^−/−^* larvae, as well as a reduction in strongly habituating ROIs in the hindbrain. Notably, this resembles the distribution of ROIs in the paradigms that drive slower habituation in WT animals (f60 and s60, Supplemental Figure 4), and provides hints that similar network changes may underlie slower habituation in both cases.

We then applied graph theory to these results (Figure 4c), looking first at correlations among 90 nodes (having eliminated nine of the original 99 nodes with a requirement that all nodes be represented in at least three larvae). Generally, correlations across the network were stronger in WT than in *fmr1^−/−^* in the first trial (resulting in positive values shown in blue). This is reversed in the 2^nd^ and 3^rd^ trials, where the WT network habituates more quickly, leaving negative (red) values that indicate persistent *fmr1^−/−^* network activity. Consistent with behavioral data, the overall correlations across the WT and *fmr1^−/−^* networks are equivalent by the 10^th^ trial, but WT networks are stronger across the core perceptual pathway (tectum, thalamus, and pallium) described above, while *fmr1^−/−^* correlations are stronger across edges that habituate quickly in WT. Again echoing free-swimming behavior, the *fmr1^−/−^* animals show dramatically broader and stronger network correlation in the 11^th^ trial, following a break in the stimulus. All of these correlation-based observations carry through to participation across the networks (Figure 4d, e), including in the same nodes whose edges showed differences in correlation (Figure 4c).

By assessing correlation strengths across the network in a way that represents nodes’ functional and anatomical properties, we then outlined the overall functional architecture of the habituating *fmr1^−/−^* brain versus WT. First, we organized our brain-wide node-to-node relationships by functional cluster (Figure 5a, Supplemental Figure 5), allowing the level of correlation within and across clusters to be assessed. This shows that by the 2nd stimulus, there are still strong functional connections among red-red edges and along blue-blue edges in WT, and that this is largely restricted to red-red edges by the 3rd trial. By the 10^th^ trial, strong correlations only exist in red-red edges (and a few to inhibited (purple) nodes). A subset of red-blue, blue-blue, and blue-green nodes reconnect in the 11^th^ trial, reflecting recovery. In all regards, these effects resemble the habituating network dynamics shown for the f20 paradigm in Figure 3, where behavioral habituation tracks with a loss of communication between weakly habituating (red) nodes and strongly habituating (green) nodes, connected through moderately habituating (blue) nodes. By comparison, *fmr1^−/−^* animals show strong correlations between more numerous nodes in the 2nd and 3rd trials, as well as following recovery in the 11^th^ trial (Figure 5a, Supplemental Figure 5). Correlated edges are similar between the genotypes in the first (Supplemental Figure 5) and 10^th^ trials (Figure 5A), showing that the networks are closer to equivalent in the naïve state and following habituation. Consistent with the analyses in Figure 4, this suggests that uncoupling across functional clusters occurs more slowly and recovers more completely in *fmr1^−/−^* animals, providing a mechanism by which the sensorimotor transformation is slanted towards downstream network activity and behavioral responsiveness in these animals.

**Figure 5.**
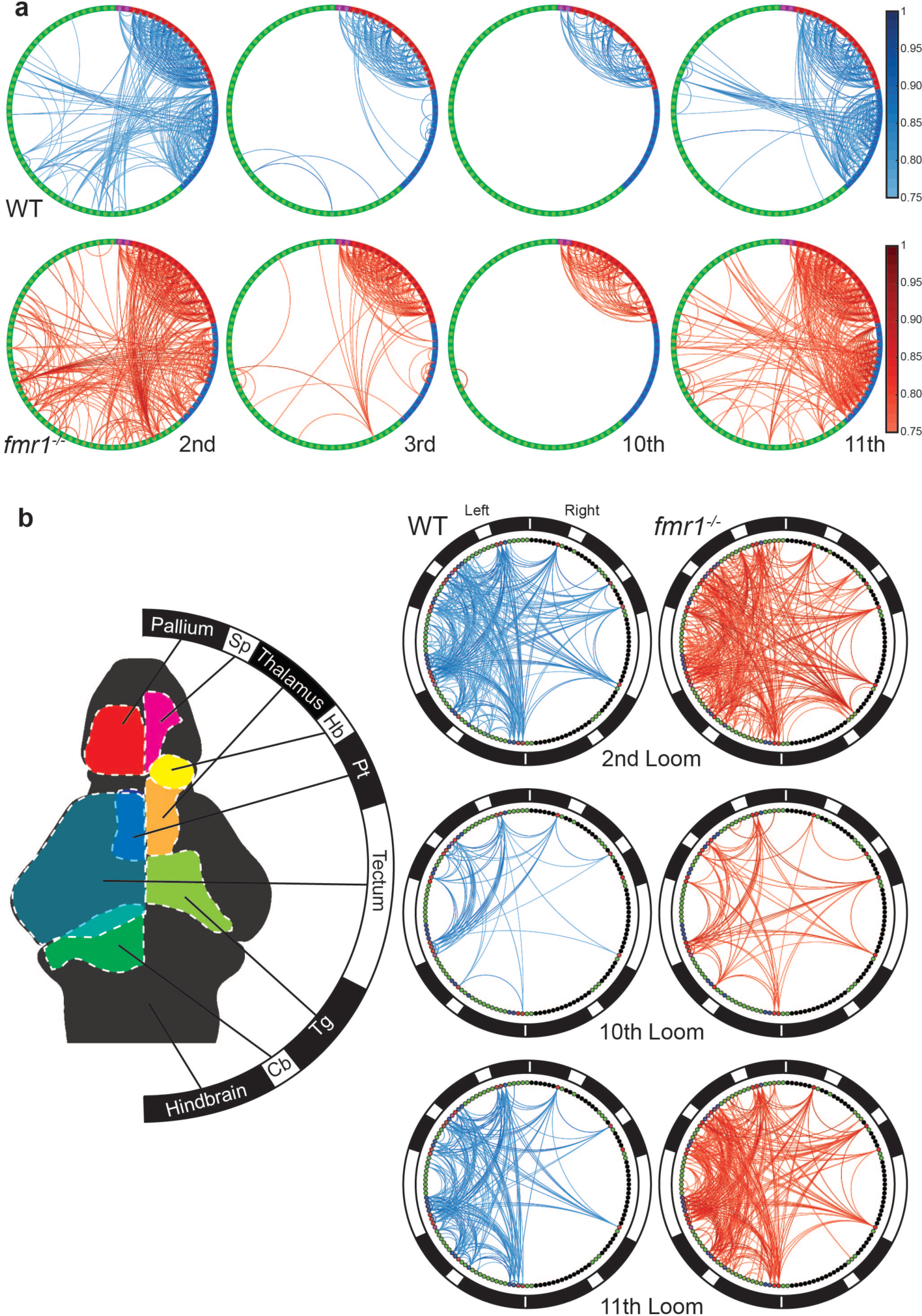
Functional and spatial habituation networks across WT and *fmr1^−/−^* brains. **a.** Edges with absolute correlation values above 0.75 are shown across 90 nodes, sorted by functional cluster (indicated by colors on the boundary). Network-wide correlations are shown for WT (top) and *fmr1^−/−^* animals (bottom) in the 2^nd^, 3^rd^, 10^th^, and 11^th^ trials. A larger version of this graph, including the brain region for each node, is included in Supplemental Figure 5. **b.** Relevant regions of the brain are shown relative to a coded boundary (left), and correlated edges between nodes (sorted by brain region) are shown for the 2^nd^, 10^th^, and 11^th^ trials (right). The black/white bands on the left are maintained to indicate brain regions on the small circles on the right. Correlations are enriched on the left side of each brain, consistent with stimulus presentation to the right eye. The networks for the first and 3^rd^ looms are also shown in Supplemental Figure 6.

To explore the spatial properties of this phenotype, we next represented these data organized by brain region (Figure 5b, Supplemental Figure 6). In WT animals, this analysis shows extensive correlation between nodes across all brain regions in the first trial (Supplemental Figure 6) that is progressively winnowed to the core perceptual circuit described above (mainly connections among the tectum, thalamus, and pallium on the side contralateral to the stimulus) as habituation proceeds. In the 2^nd^, 3^rd^, and 11^th^ trials (and to a lesser degree, the 10^th^ trial), this network is more extensive in *fmr1^−/−^* animals, showing stronger functional relationships between the tectum and other regions, and with a greater number of highly correlated edges from the hindbrain to other regions. This, in turn, echoes observations from Figure 3, which suggests that an uncoupling of spatially distinct perceptual and downstream networks drives habituation.

## A brain-wide model of visual habituation

From an anatomical perspective, the core loom perception circuit can be inferred from the edges that remain active through habituation. These include edges within and among the tectum, thalamus, and pallium (Figure 3b). The absence of habituation in these edges suggests that they are involved in perceiving a looming stimulus, and that they are upstream of the sensorimotor transformation that controls behavioral outputs. The regions most affected during habituation (especially the hindbrain, but also including a subset of ROIs in the pallium) are likely downstream of this transformation. This places the tectum at an intriguing pivot point in the overall network. It has a confirmed role as an important recipient of loom information^31-34,36,51^, and communicates in different ways with different brain regions. This includes nonhabituating correlations with the pallium and likely outputs to the hindbrain that habituate strongly (Figure 5b). Combined with the high density of moderately habituating ROIs, whose activity most closely mirrors behavioral habituation (Supplemental Figure 4c), in the tectum (Figure 2h), this raises the possibility that circuits within the tectum are responsible for the key changes in the sensorimotor transformation that produce habituation. This idea is reinforced by the drops in correlation between moderately habituating ROIs and weakly habituating ROIs (blue-red edges) and between moderately habituating and strongly habituating ROIs (blue-green edges) during habituation. This provides a mechanism by which moderately habituating neurons in the tectum could uncouple the core visual circuit of weakly habituating (red) neurons from downstream circuits as habituation proceeds. These uncoupled circuits, principally comprising strongly habituating (green) ROIs, show interesting diversity reflective of distinct impacts that novelty and saliency play in different brain regions. The hindbrain’s strongly habituating ROIs are more likely to correlate with the animal’s actual escape responses (Supplemental Figure 7), as are those in other motor-associated regions including the cerebellum, pretectum, thalamus, and tegmentum. This suggests an interaction with these regions’ premotor and motor circuits^50,51^ and an acute role in escape. Other strongly habituating ROIs that uncouple from the tectum occupy the pallium, including the Dm, a fear processing area^52,53^, and these are less likely to correlate to behavior on a trial-by-trial basis (Supplemental Figure 7), suggesting reduced higher-order representations of threat during habituation that are independent of trial-by-trial escape. The overall interpretation is that habituation involves the uncoupling of various downstream elements from visual perception circuitry, and implicates the tectum as the likely switch for this sensorimotor transformation. While *fmr1^−/−^* animals undergo habituation through similar overall mechanisms to those seen in WT animals, their network loses these correlations more slowly, and recovers them more dramatically after a rest, consistent with these animals’ behavioral phenotype (Figure 4a).

Overall, we have shown for the first time that neurons with distinct response profiles to repetitive visual stimuli are present throughout the brain, and that the detailed responses of these categories of neurons can be modulated by the saliency and temporal details of the stimuli. These response profiles, viewed brain-wide at cellular resolution, reflect the rates of behavioral habituation to repeated looms, providing a framework for understanding the brain-wide network changes that mediate habituation. Using graph theory, we have shown that behavioral habituation tracks with a functional disconnection of a principally visual circuit in the fore- and midbrain and a response circuit that includes known premotor regions located in the hindbrain and higher-order representations of threats in the forebrain. The central location of the tectum (homologous to the mammalian superior colliculus) in this functional network, and the prominence of moderately habituating tectal neurons whose activity reflects behavioral habituation rates, suggest that this region is involved in visual learning. Given these properties, it could serve as a pivot point for the sensorimotor transformation, a role that may be conserved in birds and primates^54^. We have shown that this overall network is present in *fmr1^−/−^* animals, but that its dynamics are shifted toward higher network correlations, greater transmission through the tectum, and ultimately slower behavioral habituation. This reveals a brain-wide mechanism for slower sensorimotor learning that reflects previously observed behavioral phenomena in animal models and humans with FXS. Importantly, it provides a departure point for targeted explorations of the circuit-level causes of learning and sensorimotor deficits in FXS and related psychiatric conditions.

## Methods

### Animals

All zebrafish (*Danio rerio*) work was performed in accordance with The University of Queensland Animal Welfare Unit (approval SBMS/378/16). Adults were reared and maintained in a Tecniplast zebrafish housing system under standard conditions using the rotifer polyculture method for early feeding 5 to 9 days post fertilization. For the visual habituation experiments with different stimulus trains we used *nacre* zebrafish embryos of the TL strain expressing the transgene, *elavl3:H2B-GCaMP6s^55^.* For the *fmr1* experiments, zebrafish embryos were bred by incrossing the fourth generation of zebrafish heterozygous for fmr1^hu2787^ ^56^ and *elavl3:H2B-GCaMP6s*, to produce clutches with a 1:2:1 Mendelian ratio (wild type: heterozygous: homozygous) for fmr1^hu2787^. The *fmr1*^hu2787^ mutants have a change (C to T) in the *fmr1* coding region leading to a nonsense-mediated-decay and the loss of the protein^56^. All fish were produced by natural spawning and reared in Petri dishes with embryo medium (1.37 mM NaCl, 53.65 µM KCl, 2.54 µM Na_2_HPO_4_, 4.41 µM KH_2_PO_4_, 0.13 mM CaCl_2_, 0.16 mM MgSO_4_, and 0.43 mM NaHCO_3_ at pH 7.2) at 28.5 °C on a 14-hour light: 10-hour dark cycle. After the *fmr1* experiments, larvae were genotyped as previously described^57^.

### Stimulus train for behavioural experiments

The stimulus train consisted of three blocks of 10 looms with 5min of rest (with a white screen) between each block. The loom was initiated with a dot that started expanding after 1s. The minimum angle of the loom was ~11° and the maximum angle of the loom was ~90°. The fast looms reached their maximum angle in 2s and the slow looms in 4s. This was followed by 2 seconds of black screen and a 9s slow fade back to white, designed to avoid any neural OFF responses. The screen remained white until the next loom initiation for a variable duration depending on the desired inter stimulus intervals (ISI) of 18, 20, or 22s for the f20 and s20 paradigms and 54, 60, or 66s for f60 and s60. A sound stimulus of 300Hz at ~85 dB was played 3 times for 1s with 1s ISI. The first presentation was 25s before the 21^st^ loom. The video and sound were displayed by a monitor (10.1 1366×768 Display IPS + Speakers - HDMI/VGA/NTSC/PAL, Little Bird, Australia). Since the sound stimulus did not produce any marked dishabituation, and was not confirmed acoustically within the chamber, we did not analyse this aspect of the experiment.

### Behavioural experiments

Individual 6dpf larvae were placed in each well of the 12 well arena (circular plugs of agar were removed to produce the wells). The wells were filled with embryo medium and were placed at 1cm above a screen inside a dark chamber, and all larvae received the same stimulus train. The chamber was kept in the dark but was illuminated with infrared LEDs. A Basler acA1920 camera recorded the movements from above, a lens (40mm Thorlabs) and a 665nm longpass filter (FGL665 - Ø25 mm RG665 Colored Glass Filter, Thorlabs) delivered infrared light to the camera with a weak signal from the screen that confirmed the timing of the looming stimuli. Movements were tracked in bins of 1s using the zebrafish tracking Viewpoint software (ZebraLab, ViewPoint Life Sciences, France), tracking three speed categories: <0.5mm/s, 0.5-30mm/s, and >30mm/s. The output of the tracking was then analysed using a Matlab script. Escape responses were defined as one or more movements above 30mm/s during a loom presentation. Further statistical analysis and graphs were made in GraphPad Prism v7.04 and R 3.5.1 (R core team, 2018). The sound failed to produce a clear dishabituation so this effect was not further analysed. The fitted curves were done in GraphPad Prism v7.04 with the exponential one phase decay curve from the 1^st^ to the 10^th^ loom of each block, using a Least Squares regression and plateau to 0. To produce the GLMM, we used the lme4 and MuMIn R packages to generate the GLMM and to calculate the R^2^. The model was fitted for a binomial distribution with the formula: response= loom+speed+ISI+ (1/fishID).

For the *fmr1* experiments the procedures were the same, however the stimulus train was a shorter version of the s20 with 20 looms instead of 30, as we did not observe an effect of the auditory tone between the 20^th^ and the 21^st^ loom. When the experiment ended, larvae were processed for genotyping. The quantification of the data was performed blind to the genotype for the *fmr1* experiments. The binomial test was performed one sided with the escape responses of the *fmr1* or Het larvae in each loom versus the probability of response of the WT for that same loom.

### Sample preparation for calcium imaging

Imaging was performed on 6dpf larvae that were embedded upright in 2% low melting point agarose (Sigma, A9045) and transferred to a 3D printed imaging chamber^58^. Imaging chambers were filled with embryo medium once the agarose had set and the tail was freed^59^ so that escape responses could be monitored. The imaging chamber was composed of a 3D-printed base (24 × 24 mm) with four posts (3 × 3 × 20 mm) raised along the four corners of the platform. The four outward faces of the chamber were fixed with a glass coverslip (20 × 20 mm, 0.13-0.16 mm thick). A glass window on the bottom of the chamber allowed filming of tail movements^58^. For the *fmr1* experiments, larvae were processed for genotyping when the experiment ended.

### Loom stimulus train for calcium imaging

Looms were presented on a 75 × 55 mm LCD generic PnP monitor (1024 × 768 pixels, 85 Hz, 32-bit true colour) with a NVIDIA GeForce GTX 970 graphics card. The monitor was positioned 30mm to the right of the larvae, and was covered by a coloured-glass alternative filter (Newport, 65CGA-550) with a cut-on wavelength of 550 nm. The minimum angle of the loom was ~10**°** and the maximum angle the loom covered was ~82**°**. The auditory stimulation (a 100Hz sound at 100dB before the 21^st^ loom) was presented with two audio speakers (Logitech Z213) placed at ~20cm from the fish. The background noise level was 40dB. As for the behavioural experiment, in the *fmr1* experiments, the procedures were the same but with a shorter version of the s20 stimulus train.

### Microscopy

Zebrafish larvae, individually mounted in the imaging chamber, were imaged for *elavl3:H2B-GCaMP6s* on a custom-built SPIM microscope^58,60^. To avoid stimulating the eyes with the light sheet, the side laser path of the SPIM was blocked, and the front SPIM plane was restricted to a space between the eyes using a vertical aperture. Captured images were binned 4 times to a final resolution of 640 × 540 pixels at 16-bit in tagged image file (TIFF) format. Fifty transverse sections at 5µm increments were captured and imaged at 2 Hz. Recording of the brain activity started 30s before the first stimulus onset and stopped after the return to white from the last loom of each block, resulting in three separated acquisitions. To image the larva and record its tail movements, a 4x 0.1NA Olympus microscope objective (PLN 4X) was placed below the sample chamber^61^, coupled with a tube lens projecting the image onto a Basler acA1920 camera, recording at 30 fps. At the end of each experiment, a single high-definition scan of non-binned images was recorded with 100ms exposure time and 2µm increments to be used for the registration of the brain of each fish (see below).

### Analysis of calcium imaging data

Calcium imaging data from the three acquisitions was concatenated in ImageJ v1.52c as a combined time series and then separated into individual slices (50 planes per fish). Motion correction was performed using Non-Rigid Motion Correction (NoRMCorre) algorithm^62^, and fluorescence traces were extracted and demixed from the time series using the CaImAn package (version 0.9)^63,64^

(http://github.com/flatironinstitute/CaImAn). We used 4000 components per slice to ensure that we would not miss any ROIs during the initialization step of CaImAn. The risk of over-segmentation was mitigated by a merge step using a threshold of 0.8 to merge overlapping ROIs. The order of the autoregressive model was set at 1 to account for the decay of the fluorescence, our acquisition speed being too slow to account for the rise time. The gSig (half-size of neurons) was set at 2, based on estimates of the sizes of the nuclei in our images. We did not use any temporal or spatial down-sampling and the initialization method was ‘greedy_roi’. The components were updated before and after the merge steps, empty components were discarded, and the components were ranked for fitness as previously^63^.

### Analysis of whole-brain activity data

For the experiment with four stimulus trains, the resulting ROIs and fluorescent traces from the CaImAn package were pooled from larvae of each stimulus train (n of the 4 datasets: f20=11, f60=8, s20=10, s60=10), and then z-scored per dataset. A K-means clustering by cityblock distance with 50 components and 5 replicates was done for each dataset. Clusters where manually selected based on their profile responses to the looms or sound and their presence across datasets and individual fish. This produced 7 clusters selected from the f20 and s20 datasets and were used as regressors for subsequent analysis of the four datasets: three strongly habituating, a moderately habituating, a weakly habituating, and inhibited and a sound responsive clusters. All ROIs from each of the 4 datasets were modelled by linear regression to each of these regressors. As the 60 sec ISI time series were longer, the time series were trimmed around the 30 looms to perform the linear regression. ROIs with an r^2^ value higher than 0.3 were then selected for further analysis. The selected ROIs were categorized by correlation to each of the 7 selected regressors. The auditory cluster was not analysed after this point. We confirmed that the clusters could be found in most or all larvae, but 3 fish (1 from f20 and 2 from f60) were discarded because their ROIs contribution to one of the habituating clusters was above 50% of the total number of ROIs for that cluster, so they were deemed as outliers in terms of responsiveness. To find the motor evoked calcium responses, we first used ImageJ to detect the tail movements from the behavioural imaging. We used a polygon ROI covering half of the fish tail to extract the mean grey values of the time series. Substantial tail movements produced large peaks and were flagged as movement events. We then build regressors for individual larvae inserting a stereotypical GCamp6s trace to the movement timing for each larva. Finally, we used a linear regression with the motor regressor of each larva as for the habituating clusters, and selected ROIs with an r^2^ value higher than 0.2.

For the t-SNE^65^ (Supplemental Figure 1g and Supplemental Figure 2) we used the Matlab function with a correlation based distance and the following parameters: Perplexity=100, Exaggeration=20. For further analysis, we pooled together the three strongly habituating clusters and we excluded the sound response cluster, resulting in four main clusters.

To calculate the proportions of ROIs for a given cluster that appear in each brain region (Supplemental Figure 4b), the number of ROIs of each cluster in each brain region was divided by the total number of ROIs of that cluster in the whole brain. We did this for each individual larva, created a mean for each dataset, and then averaged these values across all four datasets. For the analysis of the normalized responses in the tectum (Supplemental Figure 4c), a mean of the tectal ROIs’ response for each cluster was calculated for each individual fish, then the maximum response per loom was calculated based on the maximum z-score value in the window of the loom presentation adjusted by the baseline before each loom. These values were normalized to the first loom response, and a mean of the normalized maximum response was calculated for each dataset. To compare these tectal responses with the matching behavioral results we used the Pearson correlations coefficients.

To locate the subset of strongly habituating neurons that are involved in motor behaviours (Supplemental Figure 7), we calculated the Spearman correlation coefficient between each strongly habituating ROI from the f20 dataset and the motor regressor of its respective fish. We then selected the ROIs above a correlation coefficient of 0.3066 (the mean-0.1522-plus one SD −0.1544-). Finally, we calculated their proportion compared to the strongly habituating ROIs of each of the brain regions previously analysed.

For *fmr1* experiments, we performed a K-means with 50 components with the traces of all the fish. We then selected 8 clusters based on their possible loom responses. Then we performed a linear regression and selected the ROIs with an r^2^ value above 0.3. As their location and average calcium traces were similar to the functional clusters previously found, we classified the ROIs into the functional clusters from our original s20 dataset using correlation as described above. All data were quantified blind to genotype. For Figure 4b, we chose a random sample (n=11) of Hets to match WT (n=10) and *fmr1^−/−^* (n=11). The analysis was done using Matlab R2018b and GraphPad Prism v7.04.

### Correlation matrices and graphs

For graph theory, we simplified our system from the 144,709 responsive ROIs while preserving the functional identity and anatomical location of the responses. To do so, we performed a K-means clustering on the 3-dimensional spatial coordinates of the ROIs^66^ of each functional cluster, in each brain region, with *k* number of clusters. *k* was defined based on the number of ROIs. For regions with fewer than 200 ROIs, no node was placed; between 200-500, 1 node; between 500-1000, 2 nodes, between 1000-3000, 3 nodes; and >3000, 4 nodes. This was intended to strike a balance between including relatively sparse populations that may, nonetheless, make functional contributions, and weighting our analysis to some degree toward more abundant response types. This produced 102 nodes, but we discarded three nodes that had three or fewer fish contributing to them. For the remaining 99 nodes, we cross correlated the mean loom response of their ROIs and generated individual matrices for each larva, and each loom presentation. We then averaged the matrices of each dataset across larvae. To identify the network most similar to the 11^th^ trial of the f20 and f60 datasets, we performed a correlation between the matrices of the first 10 looms and the 11^th^ loom of the relevant dataset and identified the loom with the highest Pearson correlation coefficient.

We used the Brain Connectivity Toolbox^67^ to perform the graph analysis. We first generated weighted connectivity matrices and filtered out edges with an absolute correlation value below 0.75. We then subtracted each of the f20 loom matrices from the f60 matrices. The width and color of the edges is indicative of the subtraction weight. The participation coefficient was calculated between the four functional clusters identified previously (strongly habituating, moderately habituating, weakly habituating, and inhibited).

The *fmr1* dataset was treated similarly using the spatial nodes from the previous dataset. ROIs were assigned to each node based on the smallest Euclidian distance. By again discarding nodes with less than 3 larvae, we ended up with 90 nodes for this analysis. As before, we performed cross correlations and generated individual fish matrices for each loom presentation. The connectivity matrices were analyzed as above and the graphs were done with the subtraction of the *fmr1^−/−^* mutants from the WT.

### Registration to a reference brain

We used Advanced Normalization Tools (ANTs, https://github.com/ANTsX/ANTs) to register our results on the H2B-RFP reference of Zbrain^68–70^. The high definition stacks were used to build a common template, before registering this template to the Zbrain atlas^58^. The resulting warps were sequentially applied to the centroids of extracted ROIs to map them all in the same frame of reference. The Warped ROI coordinates were then placed in each of the 294 brain regions defined in the Zbrain atlas^70^.

### Data visualization

We used Unity to represent each ROI centroid as a sphere. Their diameter was adjusted based on the number of ROIs to be able to visualize the different clusters (Strongly habituating=2; Moderately habituating=3, Weakly habituating=4; Inhibited=6). An isosurface mesh of the zebrafish brain was generated from the Zbrain masks for the diencephalon, mesencephalon, rhombencephalon, telencephalon and eyes using ImageVis3D^71^. The mesh was imported in Unity and overlaid to the ROIs.

The colormaps used for Figures 2-5 and Supplemental Figures 5 and 6 were generated using two Matlab^®^ functions: The cbrewer function, https://au.mathworks.com/matlabcentral/fileexchange/34087-cbrewer-colorbrewer-schemes-for-matlab (Accessed in May 2019) which includes specifications and designs developed by Cynthia Brewer (http://colorbrewer.org/), and the MatPlotLib 2.0 default colormaps ported to Matlab, https://au.mathworks.com/matlabcentral/fileexchange/62729-matplotlib-2-0-colormaps-perceptually-uniform-and-beautiful (Accessed in May 2019).

The circular graphs (Figure 5 and Extended Figures 5 and 6) were made with a modified version of the code from Matlab^®^’s circularGraph toolbox. https://www.mathworks.com/matlabcentral/fileexchange/48576-circulargraph/(Accessed in May 2019).

Figures were produced using Matlab R2018b and GraphPad Prism v7.04 and assembled in Adobe Illustrator CS6.

### Data and software availability

All data and software will be made available upon request.

## Acknowledgements

We thank Ulrike Siebeck for conceptual discussion about this project, Jean Giacomotto for assistance with behavioral assays, Geoffrey J. Goodhill and Biao Sun for providing the *fmr1* mutant, and Scott lab members for feedback on the manuscript. Support was provided by an EMBO Long-Term Fellowship to G.C.V.; a fellowship from the Human Frontier Science Program (LT000146/2016) to M.A.T.; and an NHMRC Project Grant (APP1066887), ARC Future Fellowship (FT110100887), a Simons Foundation Pilot Award (399432), a Simons Foundation Research Award (625793), and two ARC Discovery Project Grants (DP140102036 & DP110103612) to E.K.S. Support was also provided by the Australian National Fabrication Facility (ANFF), QLD node.

## Author Contributions

E.M-L. contributed to the conceptual design of the project, collected and analyzed most data, and contributed to writing the manuscript. L.C. collected *fmr1* data and performed genotyping. M.P. collected behavioral data for the *fmr1* experiment. I.A.F-B. and M.A.T. built and maintained the light-sheet microscope. G.C.V. designed the data analysis pipeline for calcium imaging data and performed some data analyses. E.S. contributed to the project’s concept, guided analyses, and contributed to writing the manuscript.

The authors declare no competing interests.

Correspondence and requests for materials should be addressed to Ethan Scott.

---

**Supplemental Figure 1.**
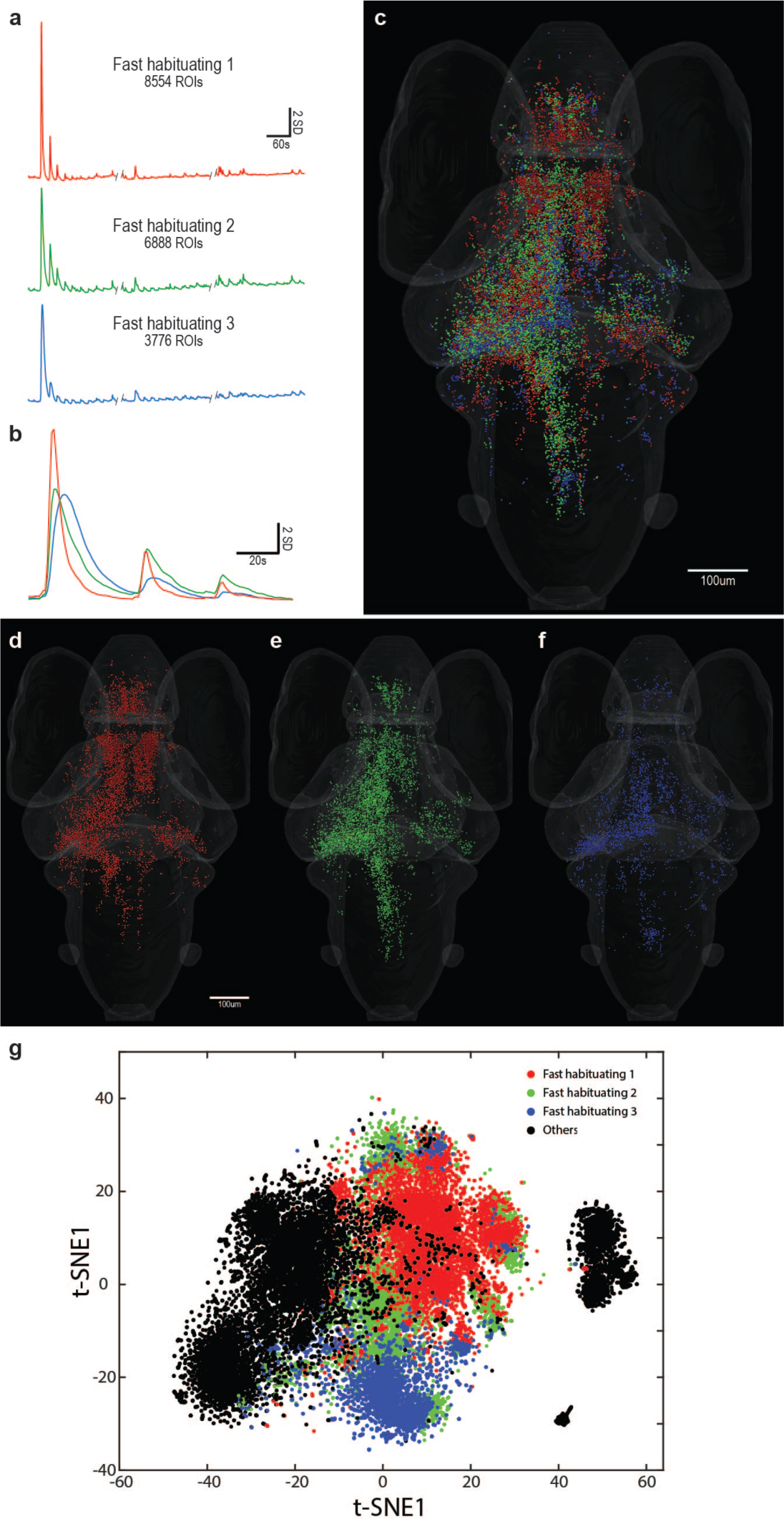
Similarities among three types of fast habituating ROI. **a.** The average responses of the ROIs composing three of the clusters produced by k-means. All are powerfully responsive the first loom stimulus, with strongly attenuated responses in the second and third trials (using the f20 stimulus train). Responses are essentially absent from trials 4-30 across three blocks of ten stimuli each. **b.** The response strengths and temporal dynamics are similar across these three groups during the first three trials, although fast habituating cluster #1 shows the sharpest response profile. **c.** The anatomical locations of these ROIs are indicated, showing a high degree of overlap in their distributions (with clusters shown individually in **d-f**). Finally, a t-SNE analysis (**g**) fails to reveal clear functional distinctions among these groups, with extensive overlap and intermingling across these three clusters, especially for the Fast Habituating 2 cluster (green). These functional and anatomical analyses form the basis for our pooling these three groups into a single “fast habituating” cluster in our subsequent analyses.

**Supplemental Figure 2.**
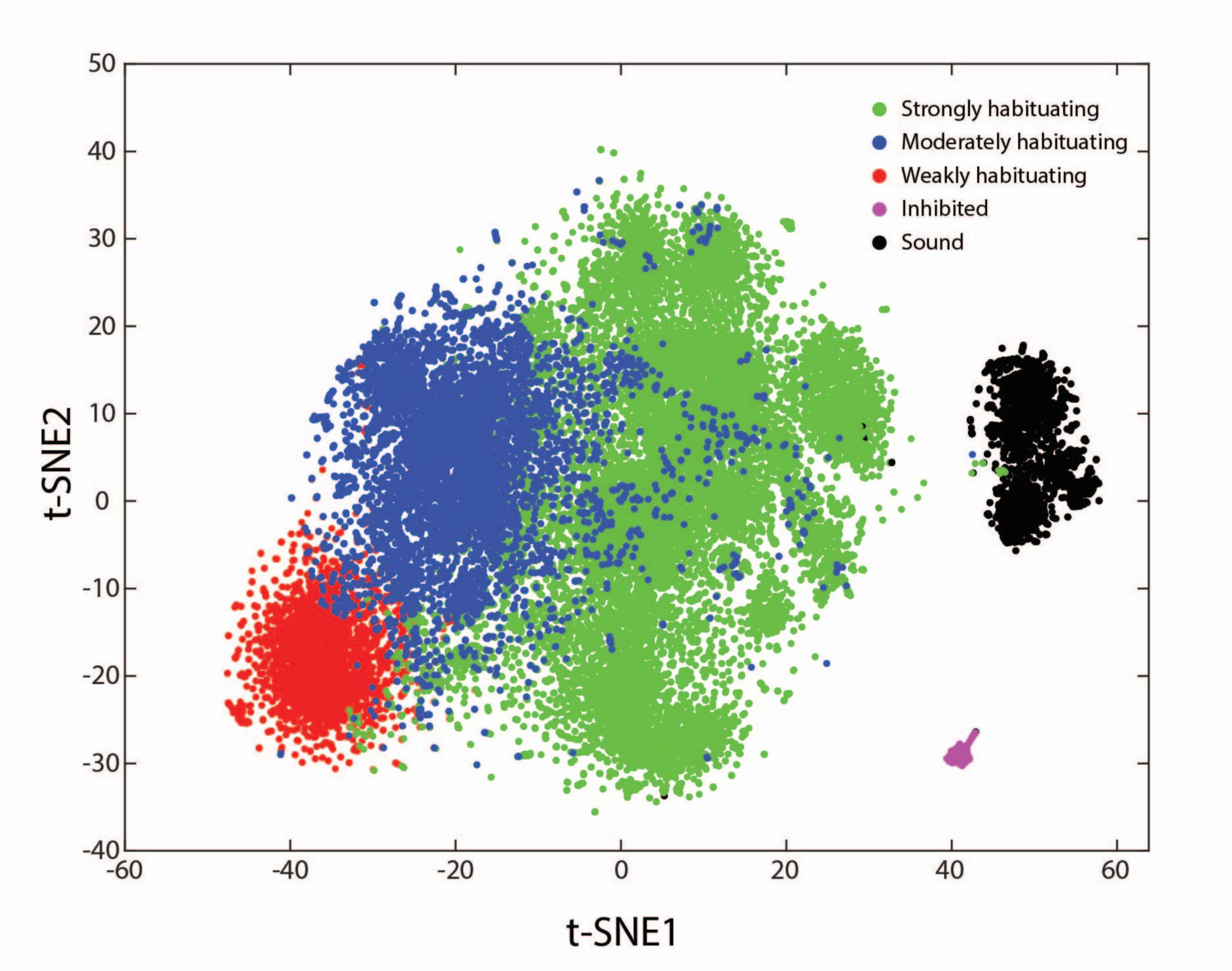
T-SNE analysis of five functional clusters. The calculated distributions of ROIs belonging to five functional clusters are represented following a t-SNE analysis. The motor associated cluster was not included because the ROIs in this cluster show different patterns of activity in different fish.

**Supplemental Figure 3.**
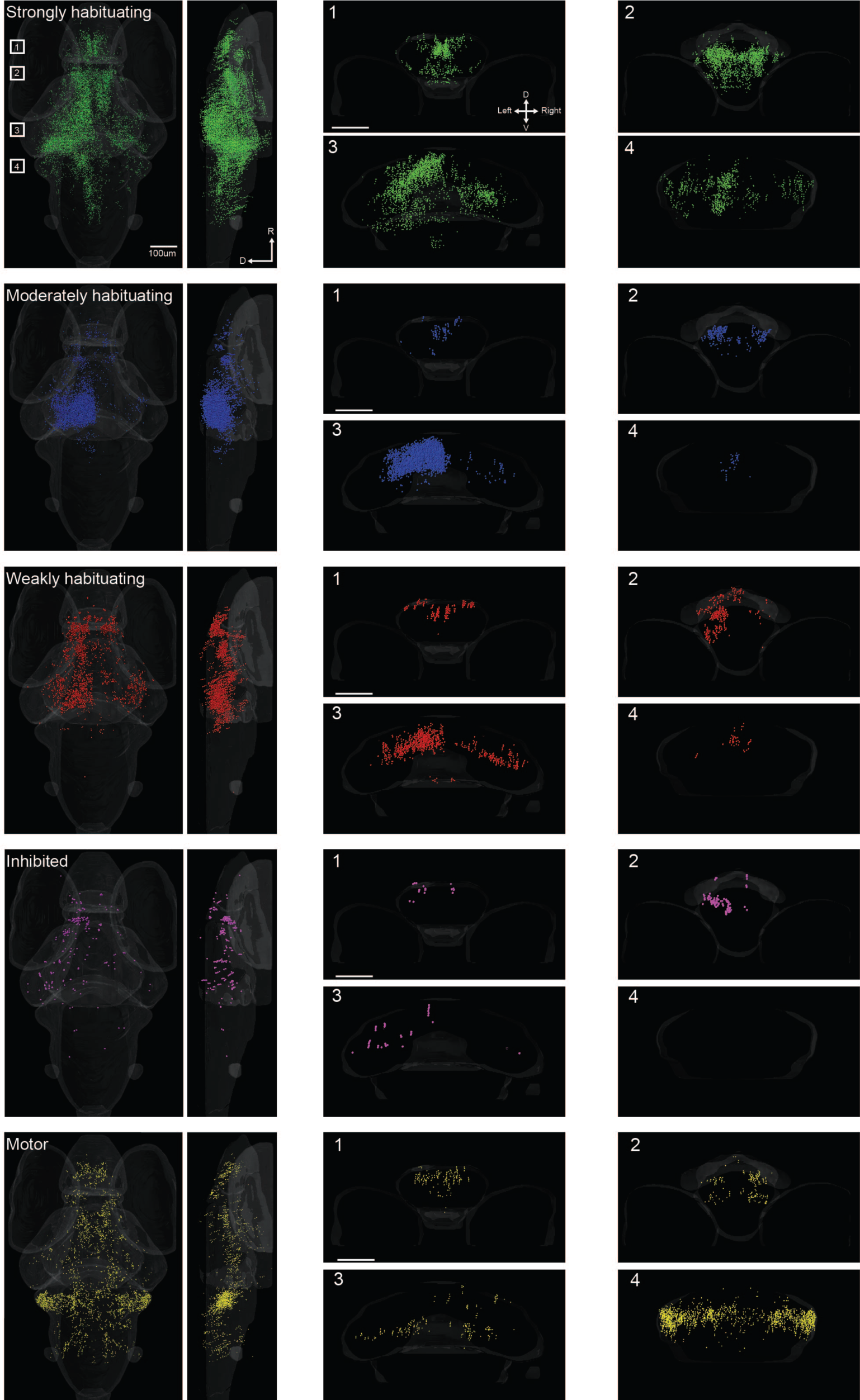
Anatomical distributions of five habituating clusters. For each functional cluster shown in Figure 2, a dorsal view, lateral view, and four coronal virtual sections are shown. The ranges included in the coronal sections are indicated in the Strongly habituating images.

**Supplemental Figure 4:**
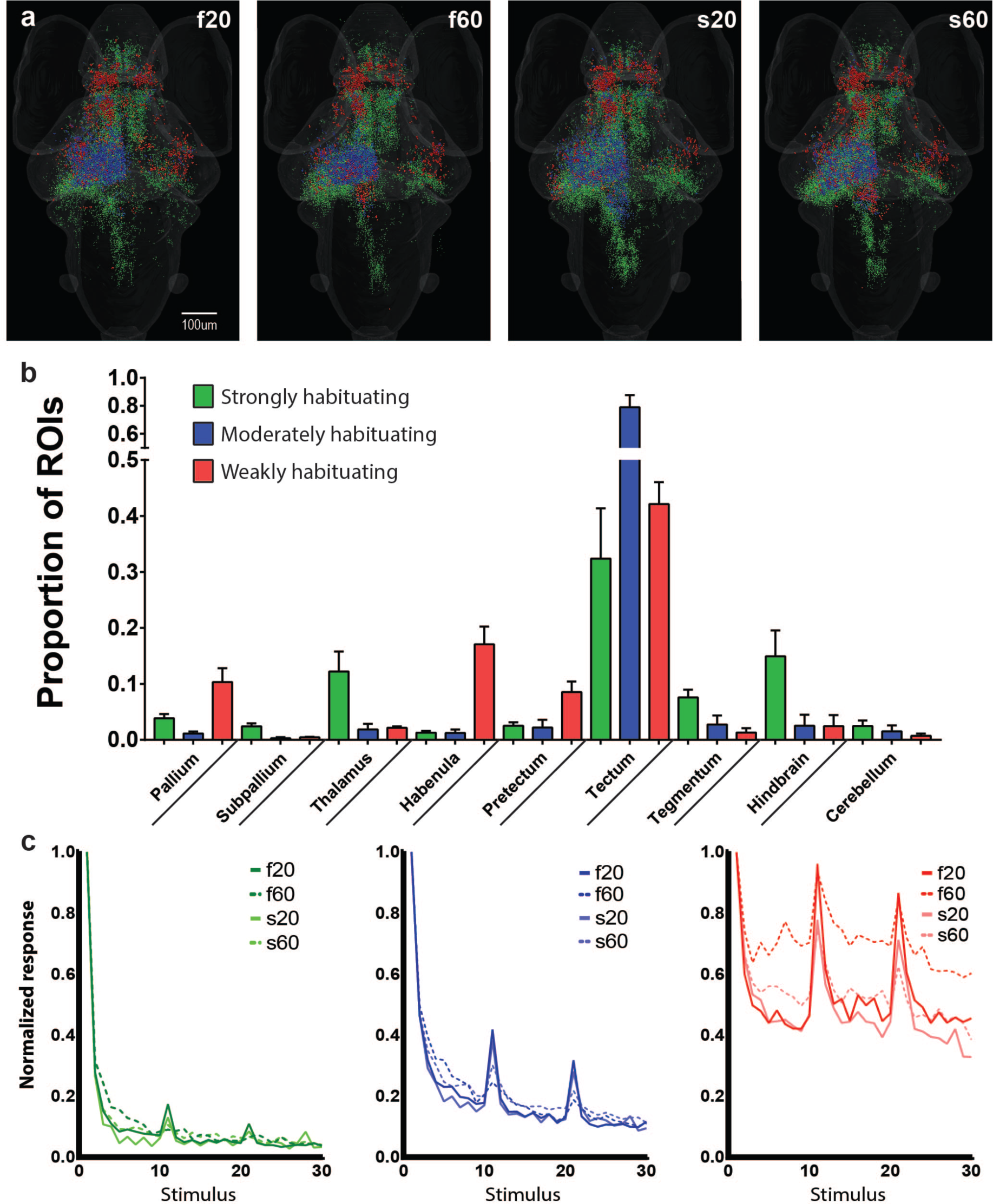
Brain-wide responses during different loom stimulus trains. **a**. Brain-wide distributions of the functional clusters from Figure 2 for each of four loom habituation paradigms. **b**. The proportions of ROIs from each functional cluster located in the indicated brain regions. **c**. The average response profiles for fast habituating (left) moderately habituating (center), and weakly habituating (right) tectal ROIs in each of the habituation paradigms.

**Supplemental Figure 5.**
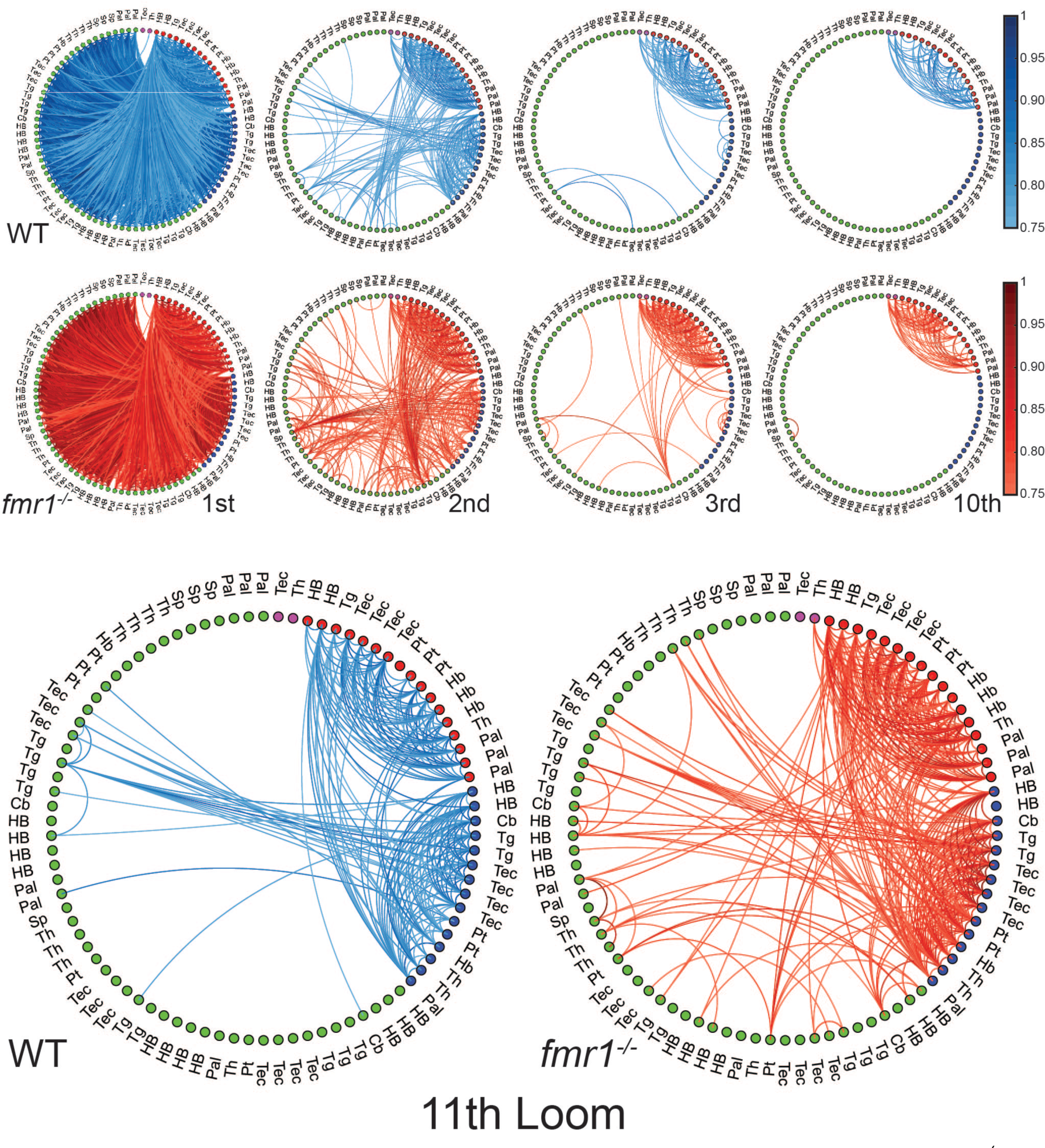
Functionally sorted brain-wide networks for WT and *fmr1^−/−^*larvae. Edges with correlations above 0.75 are shown between all combinations of nodes, and nodes are arranged by their functional clusters (colors of nodes). Networks are shown for trials 1, 2, 3, and 10 (top), and trial 11 (bottom). The brain region to which each node belongs is indicated. Pallium, Pal; subpallium, Sp; thalamus, Th; habenula, Hb; pretectum, Pt; tectum, Tec; tegmentum, Tg; cerebellum, Cb; and hindbrain, HB.

**Supplemental Figure 6.**
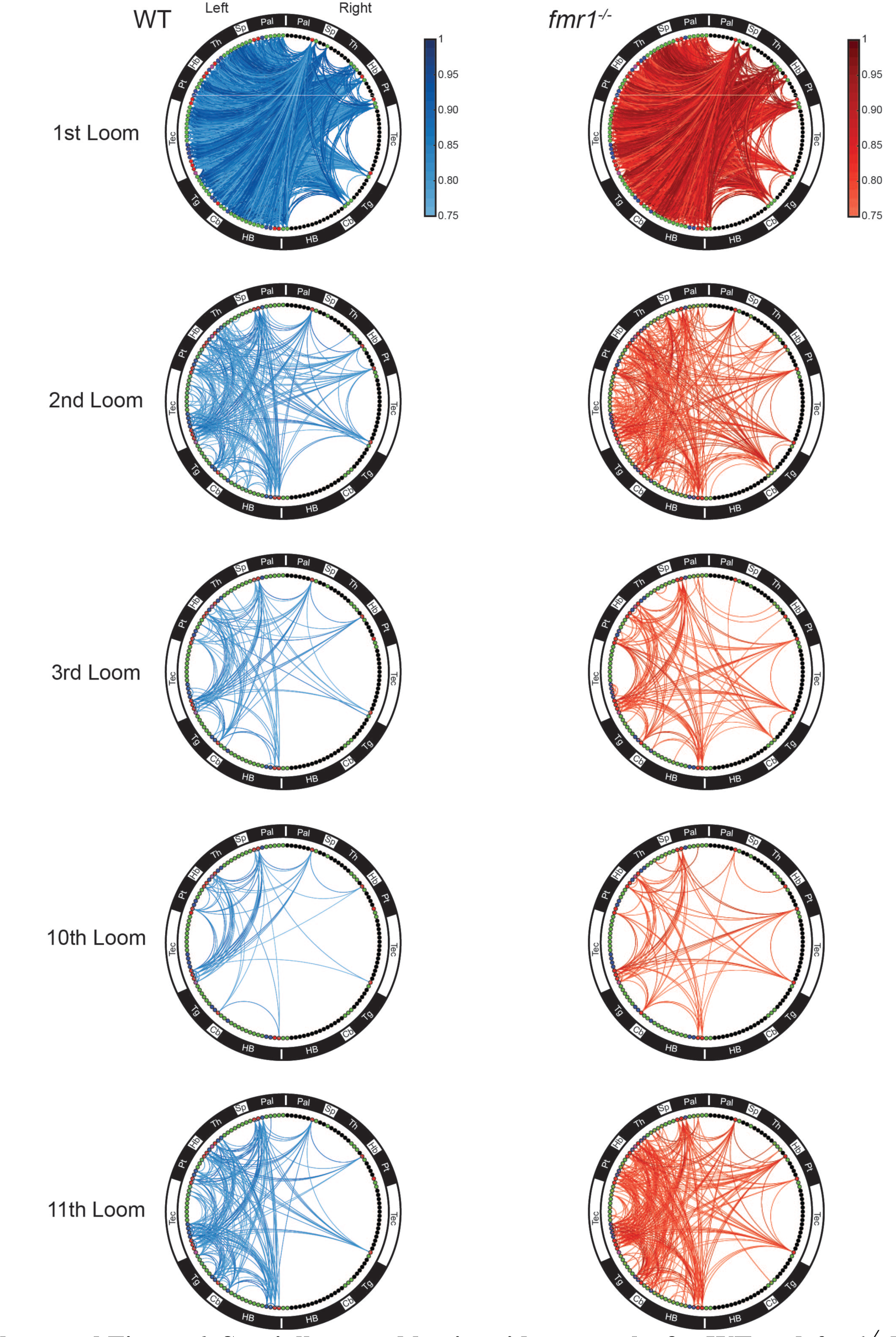
Spatially sorted brain-wide networks for WT and *fmr1^−/−^* larvae. Edges with correlations above 0.75 are shown between all combinations of nodes for trials 1, 2, 3, 10, and 11, and nodes are arranged by brain region. The nodes’ functional clusters are identified by color. Empty nodes (black) are added to match the right side (ipsilateral to the visual stimulus) to the left side spatially, despite having fewer nodes. Abbreviations are the same as in Supplemental Figure 5.

**Supplemental Figure 7:**
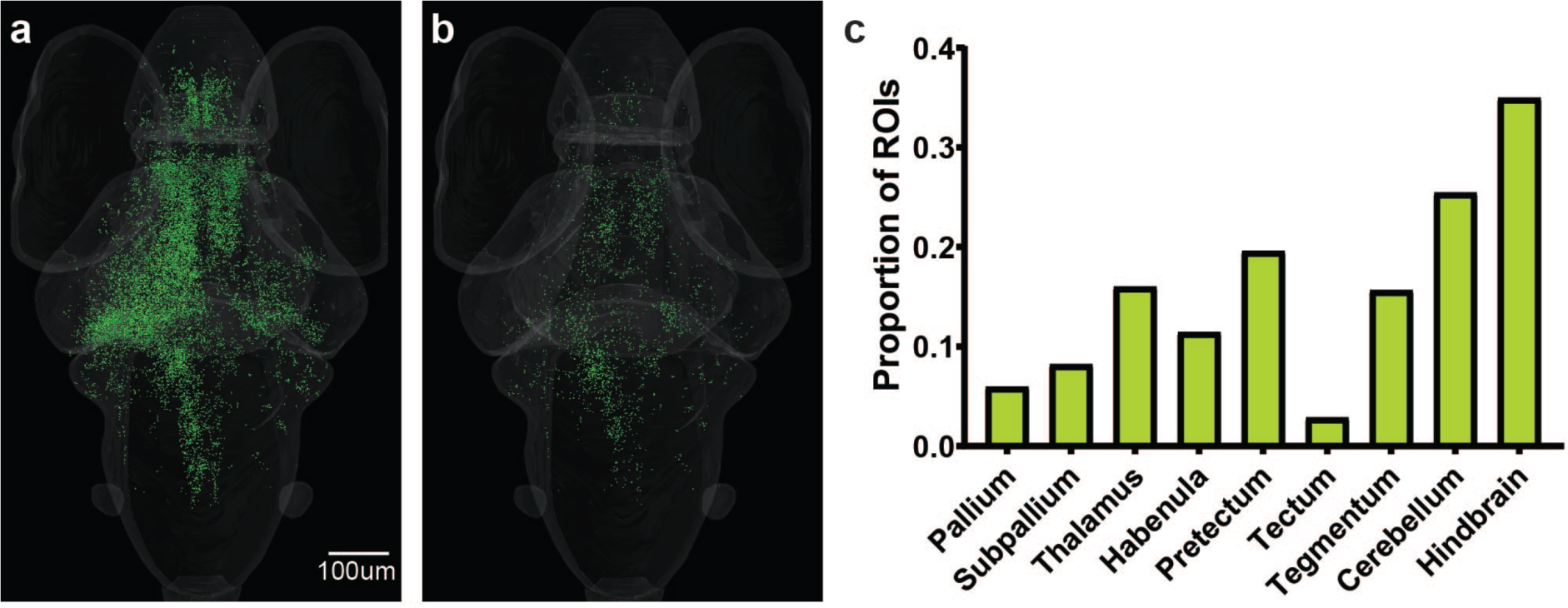
Prominence of motor-correlated strongly habituating ROIs in the hindbrain. **a.** The distribution of all strongly habituating neurons across the brain for the f20 stimulus train. **b.** The subset of the ROIs from **(a)** that show >1s.d. correlation with motor responses during loom stimuli on a trial-by-trial basis. **c.** The proportion of strongly habituating ROIs that shows this motor correlation, by brain region.

